# New estimates of the Zika virus epidemic attack rate in Northeastern Brazil from 2015 to 2016: A modelling analysis based on Guillain-Barré Syndrome (GBS) surveillance data

**DOI:** 10.1101/657015

**Authors:** Daihai He, Shi Zhao, Qianying Lin, Salihu S. Musa, Lewi Stone

**Affiliations:** Department of Applied Mathematics, Hong Kong Polytechnic University, Hong Kong, China; School of Nursing, Hong Kong Polytechnic University, Hong Kong, China; Mathematical Science, School of Science, RMIT University, Melbourne, Vic., Australia; Biomathematics Unit, School of Zoology, Faculty of Life Sciences, Tel Aviv University, Tel Aviv, Israel

**Keywords:** Zika virus, Guillain-Barré syndrome, Mathematical modelling, Infection attack rate, reproduction number, Brazil

## Abstract

**Background:** Between January 2015 and August 2016, two epidemic waves of Zika virus (ZIKV) disease swept the Northeastern region of Brazil. As a result, two waves of Guillain-Barré Syndrome (GBS), were observed concurrently. The mandatory reporting of ZIKV disease began region-wide in February 2016, and it is believed that ZIKV cases were significantly under-reported before that. The changing reporting rate has made it difficult to estimate the ZIKV infection attack rate, and studies in the literature vary widely from 17% to > 50%. The same applies for other key epidemiological parameters. In contrast, the diagnosis and reporting of GBS cases were reasonably reliable given the severity and easy recognition of the diseases symptoms. In this paper, we aim to estimate the real number of ZIKV cases (i.e., the infection attack rate), and their dynamics in time, by scaling up from GBS surveillance data in NE Brazil.

**Methodology:** A mathematical compartmental model is constructed that makes it possible to infer the true epidemic dynamics of ZIKV cases based on surveillance data of excess GBS cases. The model includes the possibility that asymptomatic ZIKV cases are infectious. The model is fitted to the GBS surveillance data and the key epidemiological parameters are inferred by using the plug-and-play likelihood-based estimation. We make use of regional weather data to determine possible climate-driven impacts on the reproductive number ℛ_0_, and to infer the true ZIKV epidemic dynamics.

**Findings and Conclusions:** The GBS surveillance data can be used to study ZIKV epidemics and may be appropriate when ZIKV reporting rates are not well understood. The overall infection attack rate (IAR) of ZIKV is estimated to be 24.1% (95% CI: 17.1% - 29.3%) of the population. By examining various asymptomatic scenarios, the IAR is likely to be lower than 33% over the two ZIKV waves. The risk rate from symptomatic ZIKV infection to develop GBS was estimated as *ρ* = 0.0061% (95% CI: 0.0050% - 0.0086%) which is significantly less than current estimates. We found a positive association between local temperature and the basic reproduction number, ℛ_0_. Our analysis revealed that asymptomatic infections affect the estimation of ZIKV epidemics and need to also be carefully considered in related modelling studies. According to the estimated effective reproduction number and population wide susceptibility, we comment that a ZIKV outbreak would be unlikely in NE Brazil in the near future.

**Author Summary:** The mandatory reporting of Zika virus (ZIKV) disease began region-wide in February 2016, and it is believed that ZIKV cases could have been highly under-reported before that. Given the Guillain-Barré syndrome (GBS) is relatively well reported, the GBS surveillance data has the potential to act as a reasonably reliable proxy for inferring the true ZIKV epidemics. We developed a mathematical model incorporating the weather effects to study the ZIKV-GBS epidemics and estimated the key epidemiological parameters. We found the attack rate of ZIKV is likely lower than 33% over the two epidemic waves. The risk rate from symptomatic ZIKV case to develop GBS is likely 0.0061%. According to the analysis, we comment that there would be difficult for a ZIKV outbreak to appear in NE Brazil in the near future.

## 1 Introduction

The Zika virus (ZIKV) was first identified in 1947 in the Zika forest of Uganda [1], and within a few years was found spreading in human populations of Nigeria [2, 3]. Transmitted through the bites of mosquito vectors (usually of the *Aedes* genus), ZIKV is an arbovirus from the family *Flaviviridae* [4, 5]. Other transmission routes have also been found (materno-fetal, sexual transmission, and via blood transfusion) but they are less common [6, 7, 8, 9]. By the 1970s, the virus was circulating widely in West Africa, although it was considered a relatively mild human infection that generally results in only fever, rash and possibly conjunctivitis [3, 10]. By 2007, the virus had escaped Africa to the island of Yap in Micronesia where, according to some estimates, it infected up to 75% of the island population [11]. ZIKV reached Polynesia in 2013, and at least by 2015, it had invaded Brazil and then very quickly the rest of South America where it reached epidemic levels [12, 13]. Since its appearance in French Polynesia and Brazil, the virus has been associated with severe neurological disorders linked to birth defects. ZIKV infection was found to pass from mother to fetus during pregnancy with the potential to result in microcephaly which causes fetal abnormalities including possible skull collapse [5]. In addition, since 2014 ZIKV was found to be strongly associated with the Guillain-Barré syndrome (GBS) amongst a small proportion of those infected [14, 15]. GBS can result in long-term muscle weakness, pain, and in some circumstances death [16]. Although there is still no proven causal link between GBS and ZIKV disease, GBS has many times been associated with ZIKV outbreaks in many countries [15], and the empirical association is unusually strong.

While considered relatively benign for decades since 1947, ZIKV disease suddenly became a major global disease threat. A Public Health Emergency of International Concern (PHEIC) was announced by the WHO on February 01, 2016 [17], in the lead-up to the Rio Olympic Games in Brazil. But until then, because of the relatively low interest in the ZIKV, surveillance in most areas was of low quality with poor coverage and consequently a large under-reporting of cases. There was little knowledge of key parameters: for example the true attack rate, the proportion of asymptomatic cases amongst infected ZIKV cases, the reproductive number. This has led to stepped up activity in surveillance and modelling efforts in recent years. But given the poor case-data available and the lack of knowledge of a reporting rate (which changed significantly in time and location) for those infected with ZIKV, results from modelling efforts have often proved to be inconsistent. Here, we take a new approach that attempts to overcome some of the problems associated with the large uncertainties associated with the reporting of ZIKV cases. Instead, we work with time series of GBS cases which should be far more reliable. We argue that a high proportion of people infected with GBS will in fact report to the doctor. Figure 1 makes clear the strong association between ZIKV cases and GBS by plotting reported cases of both diseases on the same axes. It is clear that the dynamics of the two diseases are closely in step. The unique feature of our work is that we draw on this property and fit our model to GBS data collected during and following the period of a ZIKV outbreak. We use this to infer the true numbers, and dynamics in time, of ZIKV cases.

**Figure 1:**
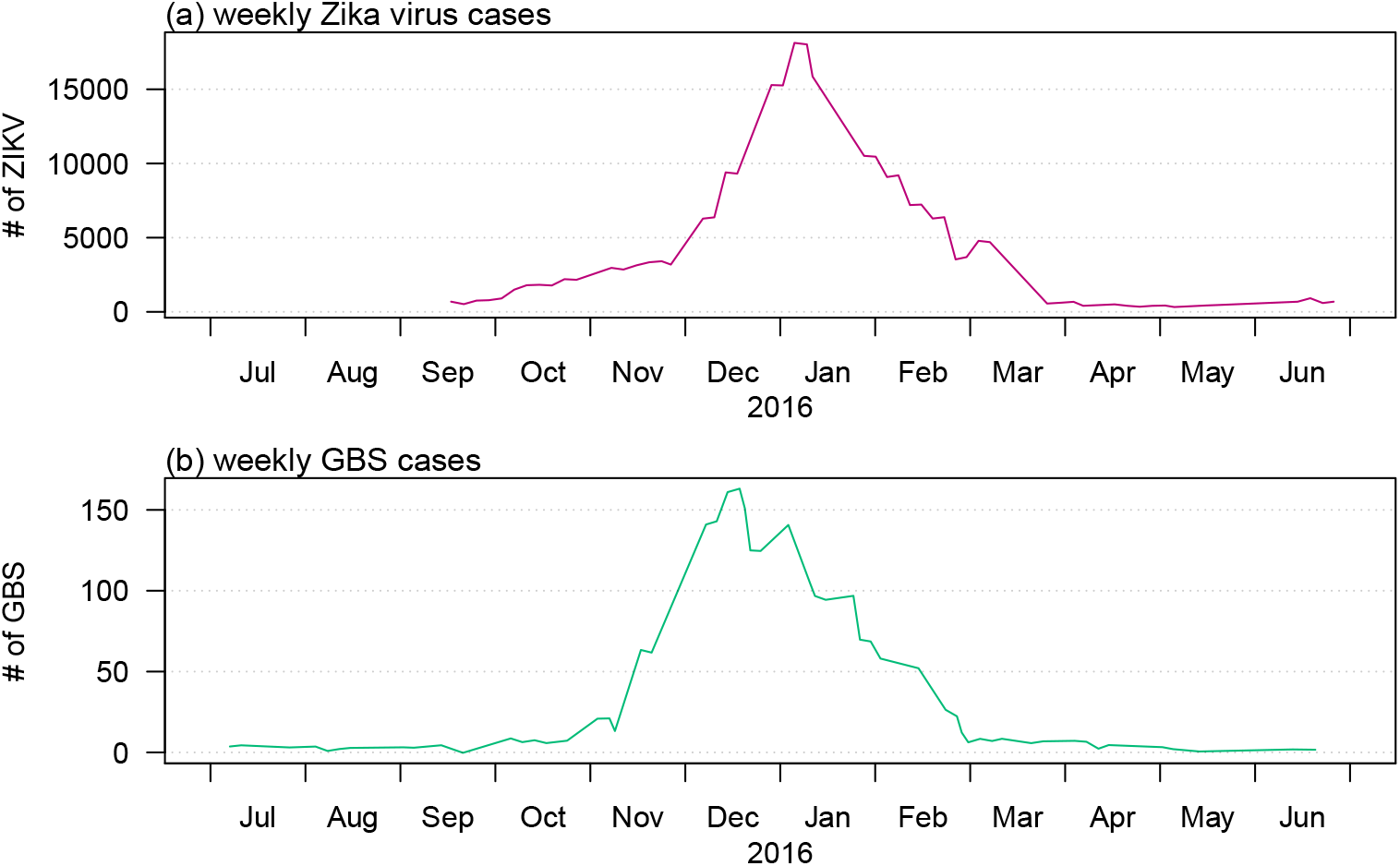
Total ZIKV (red) and GBS (green) cases as time series summed over the different states and countries: Bahia State, Colombia, the Dominican Republic, El Salvador, Honduras, Suriname, and Venezuela from April 01 of 2015 to March 31 of 2016. Data from Ref. [64].

For the modelling work that follows, it is useful to consider some of the above events in more detail on a country-specific basis, as they give further important background information that justifies our approach in using GBS as a proxy for zika-cases, on data sources and on choices of parameter values.

### French Polynesia

From October 2013 to April 2014, a severe ZIKV outbreak hit French Polynesia, and the attack rate (IAR) was first estimated as 66% [18], but updated soon after to 49% [19]. An outbreak of 42 GBS cases was simultaneously reported, but with a three-week delay in the peak timing, and was linked to the ZIKV outbreak [20]. Based on the IAR of [19], the risk of ZIKV induced GBS can thus be calculated as 0.32 GBS cases per 1,000 ZIKV infections, or just *ρ* = 0.00032. [20] estimated the proportion to be *ρ* = 0.00024. Aubry *et al.* also found that, the ratio of asymptomatic to symptomatic infections (asymptomatic ratio) was about 1:1 in the general population and 1:2 among school children [19]. These findings are notably different from estimates for a previous ZIKV outbreak in Yap island in 2007, where the asymptomatic ratio was 4.4:1 and the estimated overall ZIKV IAR was about 75% [11].

Following the ZIKV outbreak in French Polynesia, the region experienced a Chikungunya virus (CHIKV) disease outbreak with an estimated 66,000 cases from October 2014 to March 2015, and 9 GBS cases occurred [21]. The crude risk of CHIKV induced GBS was found to be 0.136 per 1,000 CHIKV infections. Thus, based on these studies [20, 21], a ZIKV infection is of (0.32÷0.136 =) 2.35-fold more likely to induce GBS when compared to a CHIKV infection. Cauchemez *et al.* [18] also found that the risk of ZIKV induced microcephaly was 95 cases (34-191) per 10,000 women infected in their first trimester during 2013-14.

### Northeastern Brazil

The Northeastern (NE) region of Brazil was the hardest-hit region in the Americas during 2015-16. In this period three mosquito-borne diseases - dengue virus, ZIKV and CHIKV, co-circulated often simultaneously, and weekly cases were documented [22]. In addition, local GBS and microcephaly cases were also recorded. Over the two years, two waves of ZIKV disease were accompanied by two waves of reported GBS cases, as shown in Fig 2, which indicated a possible epidemiological association. A striking wave of microcephaly cases with a 23-week delay to the first ZIKV wave was identified and discussed in [22]. The delay arises because ZIKV infections in the first trimester of pregnancy are most likely to induce microcephaly [18, 23, 24, 25]).

**Figure 2:**
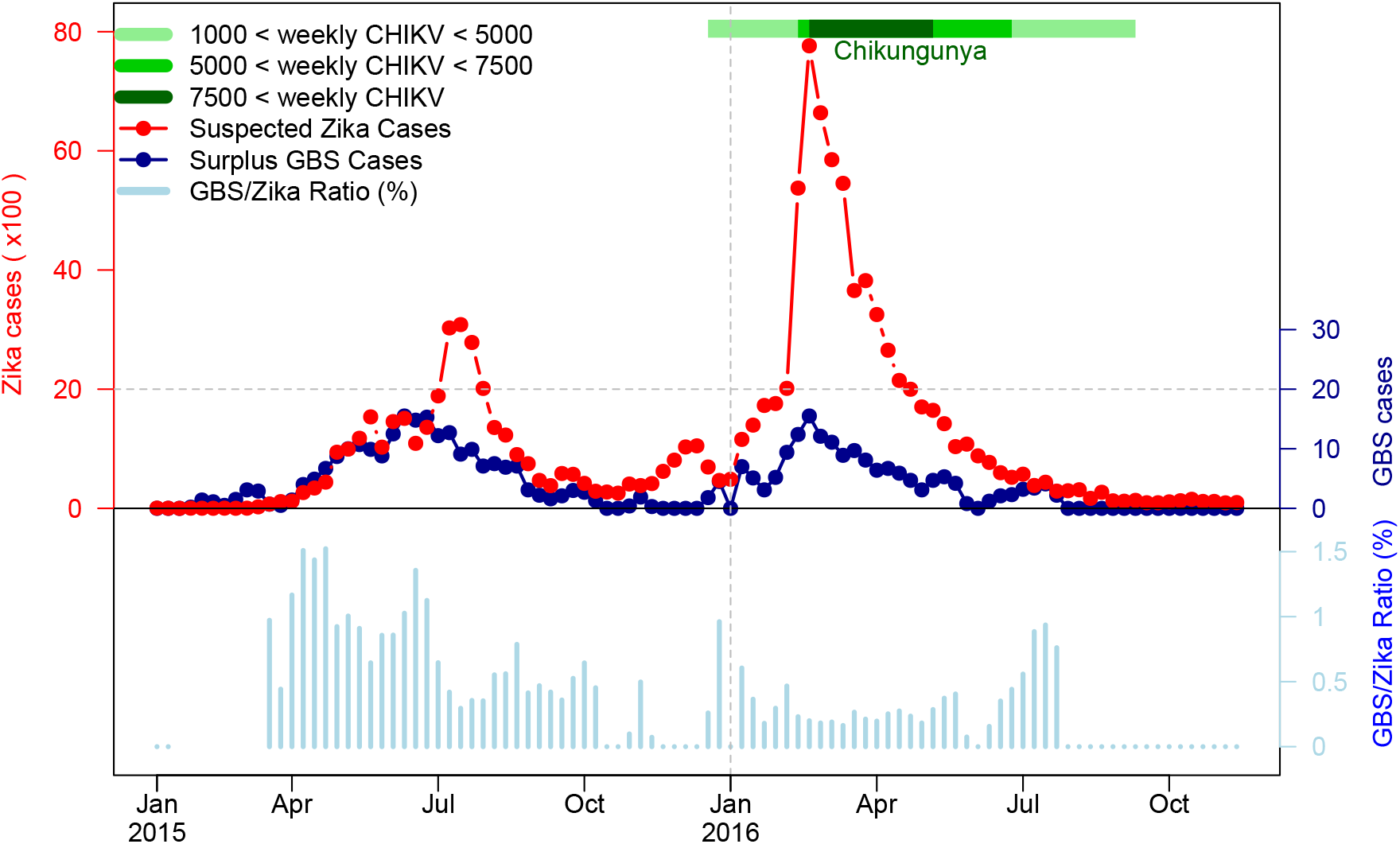
The (reported) suspected ZIKV cases, excess (or surplus) GBS cases and GBS-to-ZIKV ratio in the NE region of Brazil from January 2015 to November 2016. The red dotted line represents weekly ZIKV disease cases, the dark blue dotted line represents weekly surplus GBS cases and the light blue bars are GBS-to-ZIKV ratios. The “major” (with weekly cases over 1000) CHIKV outbreak of 2016 is shaded in green according to CHIKV disease level. The light green area denotes time periods when the weekly reported CHIKV cases were between 1000 - 5000, green denotes weekly reported CHIKV cases between 5000 - 7500 and dark green denotes weekly reported CHIKV cases over 7500. The GBS-to-ZIKV ratios are not plotted for the initial few weeks as the scale of the ZIKV data is not large enough to compute a meaningful ratio.

A substantial CHIKV wave was also observed during the second ZIKV wave in 2016 as indicated in Fig 2 and [22]. CHIKV can induce GBS with a smaller risk ratio (1 to 2.35) than ZIKV as discussed above and according to results in [21, 26, 27, 28, 29]. Note that in the latter studies, no cases of GBS induced by dengue epidemics were reported. One recent cohort study was conducted on 345 pregnant women with ZIKV rash observed (presenting at the Oswaldo Cruz Foundation) in Rio de Janeiro (the largest city in Eastern Brazil) between September 2015 and May 2016 [25]. The IAR of CHIKV was found to be approximately 17%; and in contrast, the IAR of ZIKV was 53%, as based on PCR tests. In addition, a strong cross-protection between ZIKV and CHIKV was also observed, but no cross-protection was observed between ZIKV and dengue virus (DENV). The IAR of CHIKV was 21.1%, and 41.7% for ZIKV-negative women while only 2.8% of ZIKV-positive women were infected with CHIKV. Thus, among pregnant women with rash observed in this period, the ratio of ZIKV and CHIKV is (roughly) 5 to 2. Evident cross-protection between CHIKV and ZIKV (but not between dengue and ZIKV) can be deduced from the same study with the same women [25]. Therefore, we suspect that the two waves of excess GBS cases in NE Brazil were largely due to ZIKV disease rather than CHIKV, for two reasons: (i) ZIKV is 2.35-fold likely to induce GBS than CHIKV; and (ii) ZIKV IAR could be three times higher than that of CHIKV based on the Rio de Janeiro study [25] to project the situation in NE Brazil.

Our work is based on the fact that it is difficult to estimate the infection attack rate (IAR) of ZIKV directly from the reported ZIKV cases time series given the non-constant reporting efforts over 2015 and 2016. In the literature, estimates of the IAR of ZIKV in Brazil (especially Northeast Region of Brazil) vary from less than 20% to more than 60%, and thus appear inconclusive. A summary table is provided in the Supplementary Information S4. Most of previous works were based on unreliable ZIKV surveillance data. In this work, we aim to use the relatively reliable GBS data in NE Brazil to infer the ZIKV epidemic.

The under-reporting of ZIKV cases in 2015 also appears to be reflected in what was felt to be a high number of microcephaly cases (after a 26-week delay [22]). This is because microcephaly cases are easier to identify and are thus better reported [17]. Nevertheless, the reporting criteria of microcephaly cases also changed significantly over the two years [30] leading to overall unreliable estimates. Given this known and documented unreliability [30], we felt it might not be wise to estimate IAR of ZIKV directly based on the reported number of microcephaly cases.

However, it seems a reasonable approximation to assume that the number of GBS cases per ZIKV infected individual should remain constant in time, and that the reported GBS cases are relatively well reported over time. The reporting criteria of GBS is reasonably accurate and stable owing to the distinct identifiable and severe clinical features of GBS [16]. By assuming the GBS-ZIKV risk ratio is constant, we attempted to fit an epidemic model and infer this ratio based on the GBS cases time series. Because of the co-circulation of both dengue fever and ZIKV during the two waves, misdiagnoses of ZIKV could occur [25, 23, 22], especially given both diseases have similar symptoms. Nevertheless, no GBS induced DENV was reported in the 2015 and 2016 years. Thus, the large-scale ZIKV outbreak was the major source of the excess GBS cases [22]. For these reasons, we use the excess GBS cases time series to infer the pattern of ZIKV outbreak and the overall IAR of ZIKV in Northeastern Brazil.

Mathematical modelling provides a possible way to infer the epidemic waves of ZIKV (or together with a minor proportion of CHIKV). First, we assume a constant risk ratio between symptomatic ZIKV cases and reported GBS cases (ZIKV-GBS ratio), denoted by *ρ*. Second, we simulate our ZIKV model, and fit the model to observed GBS cases with a time-dependent ZIKV transmission rate. Finally, by using iterated filtering techniques, we find the maximum likelihood estimates of *ρ* and the overall IAR.

## 2 Data and Methods

### 2.1 Data

The reported weekly excess (or surplus) GBS cases time series of NE Brazil, from Jan 2015 to Nov 2016, were kindly provided by Professor Oliveira from the Ministry of Health in Brazil, as used in their important recent study [22]. The time series are plotted in Fig 2 with datasets of ZIKV and Chikungunya for the period. The GBS data used in this work follow the case definitions given in Supplementary Information S1. In Fig 2 we observe that the GBS-to-ZIKV ratio of 2016 was significantly lower than in 2015, which was likely due to the under-reporting of ZIKV epidemic before 2016 [17].

Daily mean temperature and total rainfall (beginning from December 1, 2014) data were obtained from six cities in NE Brazil (source: https://www.worldweatheronline.com/). A map of the locations of the six cities is given the Supplementary Information S2. We calculated the daily average temperature and the average total rainfall across the six cities.

### 2.2 Methods

In previous work [6, 31], we developed a ZIKV transmission model, including both hosts and vectors, based on mosquito-borne and sexual (human-to-human) transmission of ZIKV. Hosts infected with ZIKV generate a proportion of GBS cases as determined by *ρ* which is the ratio of reported GBS cases to symptomatic ZIKV cases. In our earlier work, asymptomatic ZIKV cases were assumed to be non-infectious. However, in this work the asymptomatic ZIKV cases are now assumed to be infectious, and we study their impact on the estimation of IAR and the ratio (*ρ*). The basic reproduction number (ℛ_0_) of the model is derived and estimated. We apply the plug-and-play likelihood-based inference framework for model fitting [32].

#### 2.2.1 ZIKV-GBS Model

Fig 3 shows the model diagram of the ZIKV disease transmission pathways in both human and mosquito. Following our previous work [6, 31], we continue to assume that hosts infected with ZIKV are infectious during the convalescent stage and can infect other susceptible hosts through sexual transmission [8, 9]. However, they are assumed to be noninfectious to susceptible mosquito vectors [19, 33, 34].

**Figure 3:**
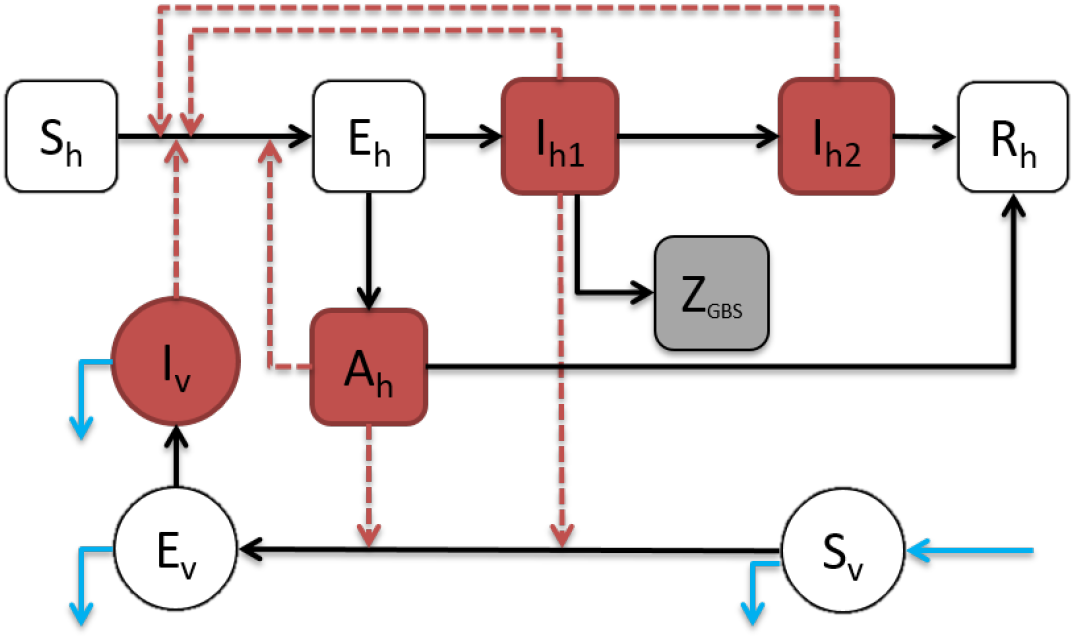
The ZIKV-GBS epidemic model diagram. The black arrows represent the infection status transition paths. Red dashed arrows represent transmission paths, and the light blue arrows represent the natural birth and death of mosquito vectors. Square compartments represent the host classes, and circular compartments represent the vector classes. Red compartments represent infectious classes, and the grey compartment is the (weekly) excess GBS cases (*Z*_GBS_). *S*_*h*_, *E*_*h*_, *I*_*h*_, *R*_*h*_ represents the numbers of Susceptible, Exposed, Infected and Recovered host population with respect to ZIKV. Please see text below Eqns (1) for complete listing of all compartment codes.

It is supposed that the asymptomatic cases are infectious at a weaker level than symptomatic cases and do not develop to the convalescent stage, which is biologically and clinically reasonable [8, 9]. We therefore arrive at the following ordinary differential equation (ODE) system (1).

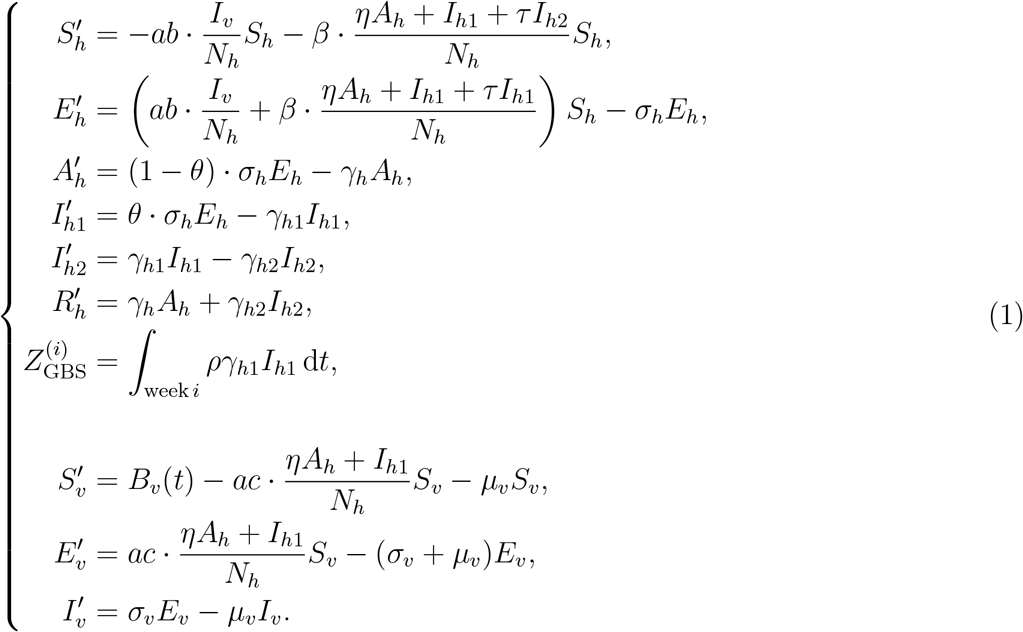

Here, *S*_*h*_ is the susceptible host class, *E*_*h*_ is the exposed host class (i.e., within ZIKV infection latent period), *A*_*h*_ denotes the asymptomatic host class, *I*_*h*1_ denotes the host class infected with ZIKV, *I*_*h*2_ denotes the convalescent host class, and *R*_*h*_ denotes the host’s recovered class. The variable 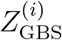 denotes the simulated weekly excess (or surplus) reported GBS cases for the *i*-th week during the study period. *S*_*v*_ is the susceptible vector class, *E*_*v*_ is the exposed vector (i.e., within ZIKV infection latent period) and *I*_*v*_ denotes the infectious vector class. The parameter *ρ* denotes the ratio of reported (excess) GBS cases per symptomatic case of ZIKV. The model (1) parameters are summarised in Table 1.

**Table 1:** Summary table of model parameters in Eqns (1). The “H” denotes human hosts’ population, and “V” denotes mosquito vectors’ population. “X→Y” denotes ZIKV infected class X infects the (ZIKV) susceptible class Y.

| Parameter | Notation | (Value)/Range | Remark/Unit | Status | Source(s) |
| --- | --- | --- | --- | --- | --- |
| Mosquito biting rate | $a$ | (0.5) 0.3 - 1.0 | per vector·day | fixed | [6, 43, 60] |
| Transmission prob. of host | $b$ | (0.4) 0.10 - 0.75 | per bite | fixed | [6, 43, 60] |
| Transmission prob. of vector | $c$ | (0.5) 0.30 - 0.75 | per bite | fixed | [61] |
| Transmission rate by contact | $\beta$ | (0.05) 0.001 - 0.10 | per day | fixed | [6] |
| Host latent period | $\sigma_h^{-1}$ | (5) 2 - 7 | days | fixed | [10, 62] |
| Vector latent period | $\sigma_v^{-1}$ | (10) 8 - 12 | days | fixed | [60, 63] |
| Asymptomatic infectious period | $\gamma_h^{-1}$ | (7) 5 - 10 | days | assumed | Nil |
| Infectious period | $\gamma_{h1}^{-1}$ | (5) 3 - 7 | days | fixed | [6, 62] |
| Convalescent infectious period | $\gamma_{h2}^{-1}$ | (25) 14 - 30 | days | fixed | [33, 34] |
| Proportion of symptomatic | $\theta$ | (50%) 20% - 80% | Nil | to be estimated | [19] |
| infectivity scale of asymptomatic | $\eta$ | 0.0 - 0.99 | H→H, H→V | to be estimated | Nil |
| infectivity scale of convalescent | $\tau$ | (0.3) 0.01 - 0.99 | H→H | fixed | [6] |
| female vector lifespan | $\mu_v^{-1}$ | (25) 4 - 35 | days | fixed | [60, 61] |
| Ratio: $\frac{\text{reported GBS}}{\text{symptomatic ZIKV}}$ | $\rho$ | 0.001% - 0.1% | Nil | to be estimated | [15, 19, 20] |
| Ratio: $\frac{\text{mosquito population}}{\text{human population}}$ | $m(t)$ | 0 - 20 | time-dependent | to be estimated | [6, 31, 43] |
| Initial susceptible proportion | $S_h(0)/N_h$ | 0.25 - 1.0 | Nil | to be estimated | [6] |

In addition,

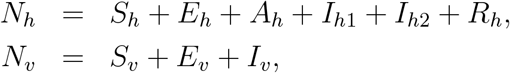

where *N*_*h*_ and *N*_*v*_ represent the total number of hosts and vectors respectively, of which *N*_*v*_ is time-dependent. The population of the Northeastern (NE) region of Brazil in 2014 was *N*_*h*_ = 56.7 million [35].

As in our previous work, it is assumed that the total mosquito population is given by:

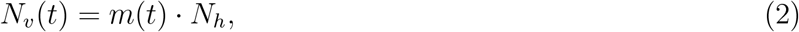

where *m*(*t*) is the (time-dependent) ratio of mosquitoes population (*N*_*v*_(*t*)) to humans population (*N*_*h*_). In the model simulation, in order to reflect the changing dynamics of *m*(*t*) to the mosquito population, we increase the susceptible mosquitoes appropriately when *m*(*t*) increases, and remove the susceptible and infectious mosquitoes when *m*(*t*) decreases to compensate. In other words, the human population (*N*_*h*_) is fixed to be constant, whereas we vary the mosquito population (*N*_*v*_(*t*)) to reconstruct the time-dependent *m*(*t*).

#### 2.2.2 Basic Reproduction Number

Following previous studies, the basic reproduction number, ℛ_0_, is derived using the next generation matrix method [6, 36, 37, 38]. We have

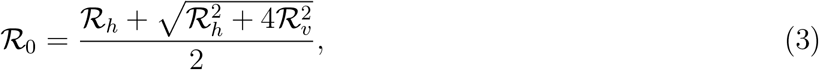

where

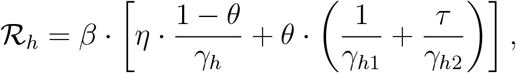

and

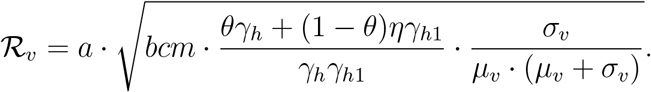

From Eqn (3), it can be seen that ℛ_0_ depends on the mosquito-borne transmission path (in term of ℛ_*v*_) and the human-to-human transmission path (in term of ℛ_*h*_). Furthermore, if one excludes the exposed and asymptomatic compartments, 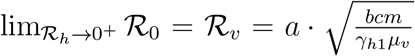, which provides the basic reproduction number of the classical Ross-Macdonald malaria model [6, 39, 40].

#### 2.2.3 Model Fitting and Parameter Estimation

To evaluate our methodology, model (1) was set up to fit the real epidemic data in NE Brazil. The time series of the number of weekly excess GBS cases in NE Brazil is modelled as a partially observed Markov process (POMP, also know as hidden Markov model) with a “spillover” rate (*ρ*) from local symptomatic ZIKV cases. Here *ρ* is the combined effect of the GBS reporting ratio and the risk rate of “symptomatic ZIKV inducing GBS i.e., the ratio 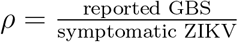 (see Table 1).

The simulated (weekly) number of excess GBS cases (*Z*_GBS_) from model (1) is considered as the theoretical or true number of cases. And the corresponding observed GBS cases of the *i*-th week, 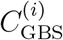, are assumed to have a Negative-Binomial (NB) distribution [6, 32, 41, 42, 43, 44].

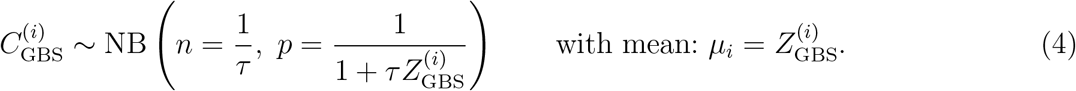

Here, *τ* denotes an over-dispersion parameter that needs to be estimated. Finally, the overall log-likelihood function, *ℓ*, is given by

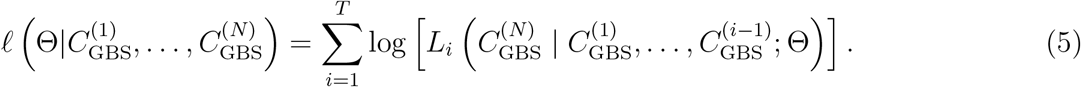

The vector Θ denotes the parameter vector under estimation. The *L*_*i*_(·) is the likelihood function associated with the *i*-th NB prior defined in Eqn (4). The term *T* denotes the total number of weeks during the study period.

Our methodology reconstructs the mosquito abundance *m* = *m*(*t*) which is otherwise unknown but variable and time-dependent over the study period. Following Eqn (3), the basic reproduction number is a function of *m*(*t*), and thus we also allow ℛ_0_ to be time-dependent (i.e., ℛ_0_ = ℛ_0_(*t*)). The time-dependent *m*(*t*) is climate-driven and modelled as an exponential function of the daily average temperature and rainfall time series, together with a two-piece step function for the baseline component. It is modelled as follows

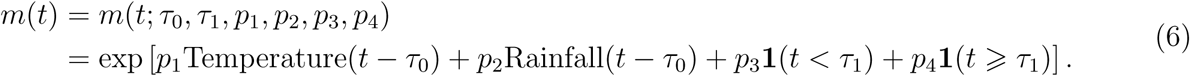

The term *τ*_0_ is the time delay between the occurrence of weather factors and their effects on the GBS epidemic. It contains the lagged effect on the local mosquito population, the progress from ZIKV to GBS development and any reporting delay. From previous studies [22, 45], there exists a time delay of at least 3 weeks between the exposure of patients to ZIKV and the development of GBS (i.e., an incubation period plus a typical reporting delay). For the mosquito, the life cycle progresses from an egg to an adult, and maturity takes approximately 8-10 days [46]. Therefore, the time lag of the effects from the weather factors are taken to be one month in total i.e., *τ*_0_ = 3 × 7 + (8 + 10)/2 = 30 days.

In Eqn (6), *p*_1_ and *p*_2_ are the scale parameters controlling the effects of local temperature and rainfall respectively. The two terms *p*_3_ and *p*_4_, are time-driven baseline effects characterizing trends in *m* that switch on depending on the time period *τ*_1_. We could view *τ*_1_ as the timing of baseline change in the mosquito population, which could be due to the interference between ZIKV and CHIKV for instance and/or local mosquito control measures. The function **1**(·) is an indicator function, which equals 1, if the condition in the brackets is true; but is 0 otherwise.

Based on fitting and comparisons, the scale of *p*_2_ was found to be negligible in magnitude, indicating that the effects of the local rainfall is (relatively) negligible, compared to temperature. Thus, in most parts of the analysis that follows, we neglect the rainfall term in Eqn (6) for simplicity.

We note that the average lifespan of the female mosquito *µ*_*v*_ is approximately 30 days. This differs from Zhang *et al.* who suggest the average lifespans goes from just under 1 day up to 7.2 days [12]. In this respect, their parametrisation seems problematic, and they probably considered the average lifespan of the mosquito, rather than the female mosquito.

According to Eqn (3), ℛ_0_ is a function of *m*(*t*), and thus ℛ_0_ is also time-dependent. Hence, ℛ_0_ can also be determined by the parameters in Eqn (6), i.e., ℛ_0_ = ℛ_0_(*m*) = ℛ_0_(*t*; *τ*_0_, *τ*_1_, *p*_1_, *p*_2_, *p*_3_, *p*_4_). Besides the climate-driven model, we also test a non-mechanistic model where the mosquito population (or transmission rate) is an exponential function of the a cubic spline function. Similar techniques were used in our previous work [43]. We compare the result with the climate-driven model and the non-mechanistic model.

The parameter fitting and inference process are rigorously and exhaustively checked within biologically and clinically reasonable ranges. We should have confidence that the fits of observed time-series are realistic because of the consistency with the true underlying epidemiological processes rather than because of artificial model over-fitting. The maximum likelihood estimate (MLE) approach is adopted for model parameter estimation. The 95% confidence intervals (CI) of parameters are estimated based on the parameter ranges in Table 1, using the method of profile likelihood confidence intervals [31, 32]. The Bayesian Information Criterion (BIC) is employed as a criterion for model comparison, and quantifies the trade-off between the goodness-of-fit of a model and its complexity [47]. The simulations were conducted by deploying the Euler-multinomial integration method with the time-step fixed to one day [32, 39]. We deploy the iterated filtering and plug-and-play likelihood-based inference frameworks to fit the reported number of excess GBS cases time series [6, 32, 43, 48, 49]. The `R` package “`POMP`” is available via [50]. Parameter estimation and statistical analysis are conducted by using `R` (version 3.3.3) [51].

## 3 Results

### 3.1 Connecting the GBS and ZIKV data, and changing reporting rates

Figure 1 plots the time series of ZIKV cases and GBS from the period April 1 of 2015 to March 31 of 2016. The data are an aggregation of the six countries Columbia, the Dominican Republic, El Salvador, Honduras, Suriname, and Venezuela as well as the Bahia State in Brazil. These time series demonstrate the tight connection between the reported ZIKV disease and GBS, whose case numbers closely mimic one another in time. The connection is the basis of our method for estimating ZIKV cases from GBS reports, which as we have discussed, are by their nature, reasonably reliable records.

The North East Brazil datasets are plotted in Figure 2. Here we see two epidemic outbreaks of reported ZIKV cases, where the second outbreak in 2016 is far stronger than the first in 2015. Despite this, the two waves of GBS appear similar over the two years although a close examination reveals there were fewer cases in 2016. If one ignores possible regional difference and adopts the GBS-ZIKV risk rate of 0.032% i.e., 0.32 GBS cases per 1,000 ZIKV infections (asymptomatic and symptomatic) calculated in [20], the total cases of ZIKV can be approximated according to the excess GBS cases time series (Fig 2). But this is a naive calculation and we will seek ways to improve this.

Tallying the case numbers, in 2015 there were 233 excess GBS cases and 38,641 reported ZIKV cases, but in 2016 there were 168 excess GBS cases and 70,916 reported ZIKV cases. The ratio of GBS/Zika reported cases is plotted (blue) in Fig 2, and one sees the transition from GBS/ZIKV(repoted) (= 233÷38641 = 0.60% in the first year (2015) to GBS/ZIKV(reported) =168÷70916 = 0.24% in the second year (2016).

Let us first assume that the GBS/ZIKV (reported) ratio did not change in time in any major way over the two years 2015 and 2016. Our analysis of data from the time series in Fig 2 shows that as GBS cases dropped from 233 cases in 2015, to 168 cases in 2016, i.e. by a factor of 0.72 (168/233), the number of reported ZIKV cases rose by a factor of 70, 916÷38, 641 = 1.8. The only explanation for this is that there must have been a major under-reporting of ZIKV cases in the first year of 2015 [44, 52]. This also seems reasonable since in 2015 the official WHO ZIKV reporting program had not yet been launched [17]. Suppose now the GBS/ZIKV(reported) ratio was 0.24% in both 2015 and 2016 even though we know that this could not be the case. A simple calculations shows that there should have been some 98,353 (= 233 × 70916÷168) ZIKV reported cases in 2015 rather than only the 38,641 cases that were reported in reality. Thus for the 2015 year it would appear that ZIKV was under-reported by a factor of 2.5 when compared to the ZIKV reporting rate in 2016.

### 3.2 Fitting the model to GBS data

We fit model (1) based on the reported excess GBS cases time series shown by the dark blue dotted line in Figure 2. This was repeated for different sets of baseline parameters. Several different (possible) values of *η* (asymptomatic ZIKV relative infectivity) and *θ* (proportion of symptomatic ZIKV infections) were considered. The *θ* = 0.5 simulations correspond to a 1:1 ratio of the symptomatic to asymptomatic ZIKV infection of [19]. And *θ* = 0.2 simulations correspond to the 4:1 ratio of the symptomatic to asymptomatic ZIKV infection of [11].

Fig 4 shows the fitting results with *θ* = 0.5 and *η* = 0.3. The mean GBS values for 1000 simulations are plotted (red) in time and fit the trajectory of the reported GBS cases (black line) closely. The grey shading gives the 95% credible interval (CI) of the case numbers for each day of the simulation. The models fits the data well, and all 95% CI cover the associated observation. This indicates the simulation outcomes are not statistically different to the observations, and thus our model successfully reconstructed the two waves of the ZIKV epidemic in NE Brazil. We estimate the time-dependent ℛ_0_(*t*) which ranged from 1.1 to 3.3 over the whole study period. The simulations determine the best fitting initial condition of susceptible population is *S*_*h*_(0) = 0.55. The inserted panel shows the parameter estimation of *ρ* found where the likelihood profile reaches the minimum BIC value. Namely, we fix *ρ* at 20 values over a range, fit the model (1) to the GBS data, and calculate the BIC. While the minimum is *ρ* = 0.00061, a value of *ρ* from 0.00005 to 0.0001 will yield an (almost) equivalent level of BIC given the flatness of the curve in this regime.

**Figure 4:**
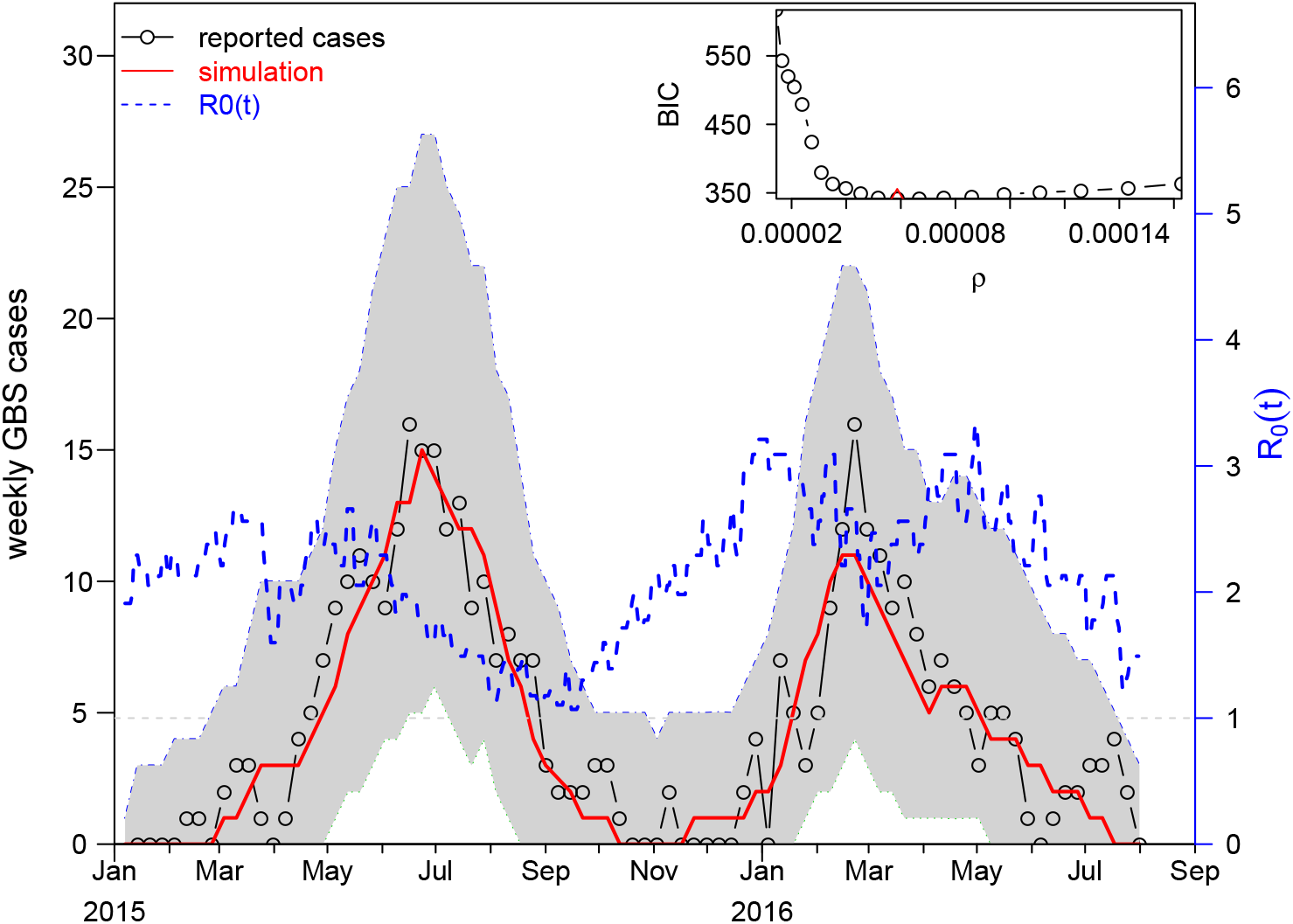
The fitting results for *θ* = 0.5 and *η* = 0.3. The fitting results in the main panel show the best scenario, which attains the smallest BIC. The red line is a plot of the mean GBS cases averaged from 1000 simulations plotted as a function of time. The grey shaded area shows the 95% credible interval (CI) of the fitted number of GBS cases. The inset panel shows the profile of BIC as a function of *ρ*. The minimum occurs at *ρ* = 0.000061, which is our best estimate for *ρ*.

In addition to the mechanistic reconstruction of ℛ_0_(*t*) in the main results here, we also present a non-mechanistic reconstruction in Supplementary Information S3. The non-mechanistic approach is implemented by using a cubic spline function to reconstruct the ℛ_0_(*t*). The model also fits the disease surveillance data well. The BIC of the non-mechanistic model is 7 units larger than the above climate-driven model in Fig 4. We find that the non-mechanistic reconstruction of ℛ_0_ matches the daily temperature reasonably well. This suggests the weather-driven ℛ_0_(*t*) in our main results here is neither coincidental nor artificial.

### 3.3 Estimation of Attack Rate (IAR) and model parameters

The estimates of the GBS/ZIKV ratio *ρ* and the IAR are summarised in Table 2. For the parameter ZIKV symptomatic ratio, *θ*, we follow the previous serological study conducted in French Polynesia that found asymptomatic: symptomatic case ratios is 1: 1 in the general population [19]. Thus, we treat the scenarios with *θ* = 0.5 in our main results. Setting a constant *θ* = 0.5, the estimation of *ρ* is roughly 0.000063 (= 0.0063%). This appears to hold even if *η*, the relative infectivity of the asymptomatics, is changed over the interval (0, 1). Estimates of *ρ* thus appear to be reasonably insensitive to the change of relative infectivity of the asymptomatics (*η*). However, *ρ* is sensitive to the change of the symptomatic proportion of ZIKV infections (*θ*). Setting *θ* = 0.2 gives *ρ* = 0.00013, but as Table 2 reveals, this result is also relatively insensitive to changes in *η*.

**Table 2:** Summary table of the estimation results of *ρ* and IAR. The estimates with *θ* = 0.5 and *η* = 0.3 are used as main results, also in Fig 4.

| $\theta$ | $\eta$ | $\rho$ | 95% CI | IAR | 95% CI |
| --- | --- | --- | --- | --- | --- |
| 0.5 | 0.1 | 0.000053 | (0.000046,0.000080) | 0.2792 | (0.1841, 0.3234) |
| 0.5 | 0.3 | 0.000061 | (0.000050,0.000086) | 0.2411 | (0.1711, 0.2932) |
| 0.5 | 0.5 | 0.000063 | (0.000049,0.000086) | 0.2352 | (0.1711, 0.3005) |
| 0.5 | 0.7 | 0.000063 | (0.000050,0.000084) | 0.2352 | (0.1753, 0.2932) |
| 0.5 | 0.9 | 0.000067 | (0.000053,0.000086) | 0.2186 | (0.1711, 0.2792) |
| 0.2 | 0.1 | 0.000139 | (0.000083,0.000169) | 0.2645 | (0.2175, 0.4423) |
| 0.2 | 0.3 | 0.000129 | (0.000117,0.000178) | 0.2847 | (0.2071, 0.3140) |

To calculate the number of ZIKV cases and IAR, we use our estimated *ρ* = 0.00061 (ratio of reported GBS to symptomatic ZIKV), and we denote our ZIKV symptomatic ratio by *θ*. The *ρ* can be estimated from the model. Then, the number of ZIKV cases equals (the number of reported GBS)÷[reported GBS/symptomatic ZIKV]÷(ZIKV symptomatic ratio), which is the number of the reported GBS*/ρ/θ*. Therefore, the IAR equals the number ZIKV cases÷the total population in the NE Brazil.

For all pairs of *θ* and *η* in Table 2, the estimated IARs are similar with IAR ≈ from 22% to 28% and the 95% CIs largely overlap. Thus, for *θ* = 0.5, we can be at least 95% sure the IAR of the ZIKV epidemic is below 33%, and is likely to be well below.

The estimates of the initial susceptible levels (*S*_*h*_(0)) and the parameters (*p*_1_, *p*_3_ and *p*_4_) that control the temporal pattern of ℛ_0_(*t*) are summarised in Table 3. Note that according to Eqn (3), *m* is proportional to 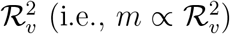, a key term in the formula for the basic reproduction number. It is not hard to show that [exp(0.5*p*_1_) - 1] × 100% is the change rate in ℛ _*v*_ when there is one unit (°C) increase in temperature. From Table 3, one unit increase in temperature will lead to an increase of (exp(0.5 × 0.52) *-* 1 =) 29.7% in ℛ_*v*_ when *η* = 0.1. And one unit increase in temperature will lead to (exp(0.5×0.53)*-*1 =) 30.3% increase in ℛ_*v*_ when *η* = 0.3. Eqn (3) shows the ℛ_0_ is comprised of ℛ_*v*_ and ℛ_*h*_, where the ℛ_*h*_ is the contribution from the sexual transmission path. The sexual transmissibility of ZIKV can be ignored owing to (i) the contribution of this path is negligibly small [6, 7]; and (ii) the sexual contact is recommended to be prevented during the ZIKV epidemics [10]. Hence, the ℛ_*h*_ could be very close to zero, and its contribution to the whole ℛ_0_ is probably far less than the mosquito-borne transmission ℛ_*v*_. According to Eqn (3), ℛ_0_ = ℛ_*v*_ when ℛ_*h*_ = 0. Provided 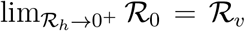, the effect of the temperature to ℛ_*v*_, determined by the *p*_1_ estimate, is (almost) equivalently applicable to ℛ_0_.

**Table 3:**
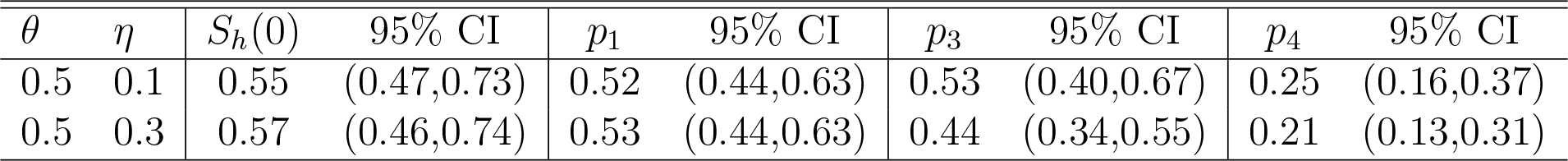
Summary table of the estimation results of the initial susceptibility (*S*_*h*_(0)) and parameters *p*_1_, *p*_3_ and *p*_4_ in Eqn (6). The estimates with *θ* = 0.5 and *η* = 0.3 are used as main results, also in Fig 4.

| $\theta$ | $\eta$ | $S_h(0)$ | 95% CI | $p_1$ | 95% CI | $p_3$ | 95% CI | $p_4$ | 95% CI |
| --- | --- | --- | --- | --- | --- | --- | --- | --- | --- |
| 0.5 | 0.1 | 0.55 | (0.47,0.73) | 0.52 | (0.44,0.63) | 0.53 | (0.40,0.67) | 0.25 | (0.16,0.37) |
| 0.5 | 0.3 | 0.57 | (0.46,0.74) | 0.53 | (0.44,0.63) | 0.44 | (0.34,0.55) | 0.21 | (0.13,0.31) |

In table 3, the *S*_*h*_(0) is estimated to be 0.55 (95% CI: 0.47-0.73) when *η* = 0.1, and 0.57 (95% CI: 0.46-0.74) when *η* = 0.3. The large overlap in the 95% CIs indicates that the two *S*_*h*_(0) estimates are not statistically different. According to the 95% CIs of *S*_*h*_(0), it is likely that over a quarter (i.e., > 25%) of the whole population were not involved in the 2015-16 ZIKV epidemic.

We estimate that the time points (*τ*_1_) when the baseline of *m*(*t*) (or ℛ_0_(*t*)) changes from *p*_3_ to *p*_4_ in Eqn (6). It was found that *τ*_1_ is most likely to be March 7 of 2016. For the parameters *p*_3_ and *p*_4_, we find significant difference in the baseline levels of *m*, which suggested the existence of the non-weather-driven temporal changes in the ZIKV transmissibility.

### 3.4 Results of the Sensitivity Analysis

As is conventional, the Partial Rank Correlation Coefficients (PRCC) are adopted to perform a sensitivity analysis of the model [6, 43, 49, 53]. Firstly, 1000 random samples are taken from uniform distributions of each model parameters. The ranges as set out in Table 1. Secondly, for every random parameter sample set, the ZVD-GBS model was simulated to obtain the target biological quantities (e.g., ℛ_0_ and total number of GBS cases in this study). Finally, PRCCs were calculated between each parameter and target biological quantities.

Results of the sensitivity analysis are presented in Fig 5, which indicates how model parameters impact the basic reproduction number (ℛ_0_) and the total reported GBS cases. ℛ_0_ is most sensitive to the vector’s biting rate (*a*), the vector to host ratio (*m*) and the vectors’ lifespan (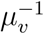, or vectors’ natural death rate, *µ*_*v*_), indicating the importance of the mosquitoes role in disease transmission. The total reported GBS cases are considerably sensitive to the proportion of symptomatic cases (*θ*), and the ratio (or risk) of excess GBS cases to symptomatic ZIKV infections (*ρ*).

**Figure 5:**
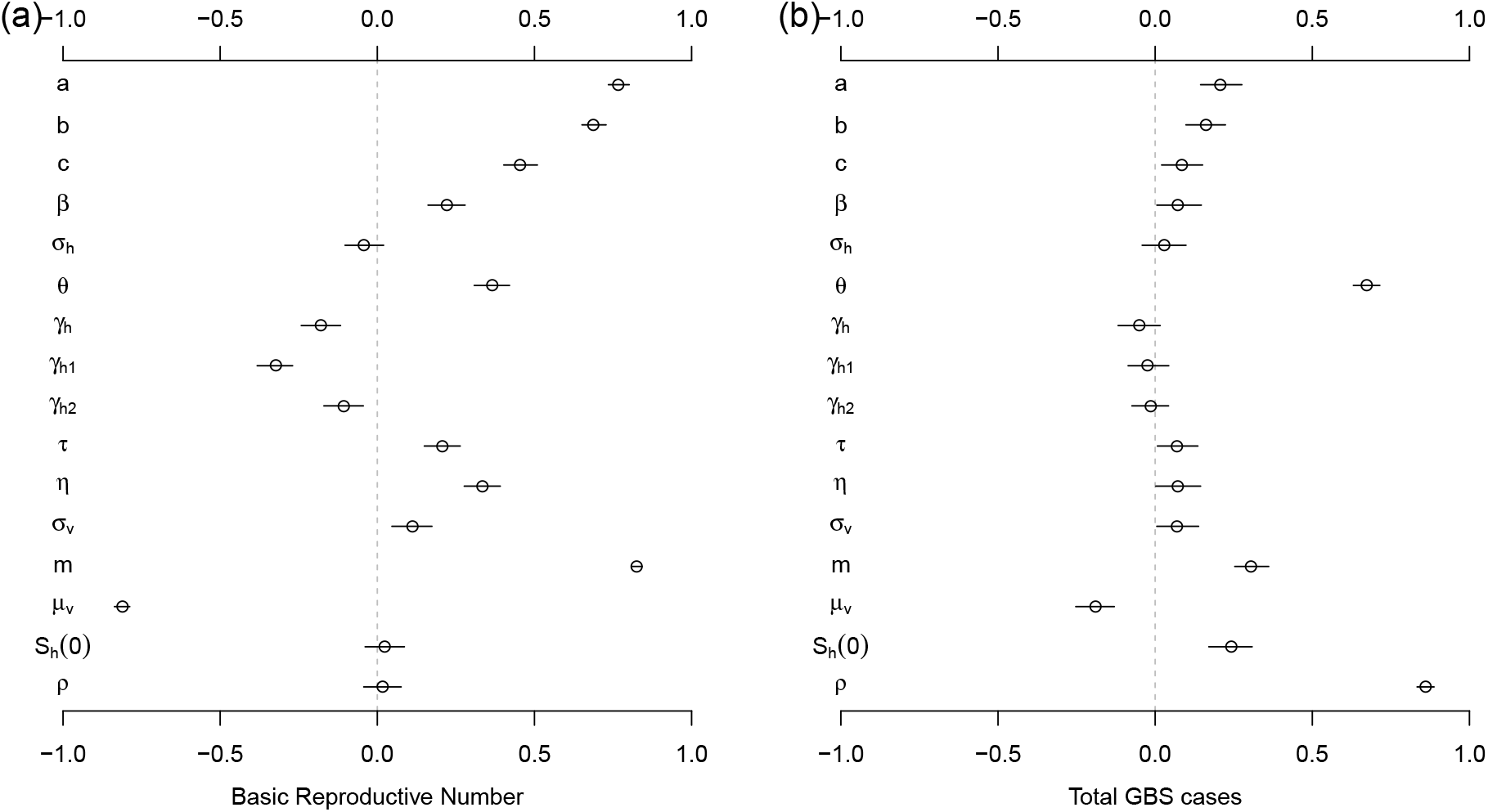
The Partial Rank Correlation Coefficients (PRCC) of the basic reproduction number, ℛ_0_, (panel (a)) and total GBS cases (panel (b)) with respect to model parameters. The *S*_*h*_(0) in this figure denotes the initial susceptible ratio, i.e, *S*_*h*_(0)*/N*_*h*_. The black circle is the estimated correlation, and the bar represents 95% CI. The ranges of parameters are in Table 1.

## 4 Discussion

Based on the striking parallel between cases of ZIKV disease and cases of GBS, as seen in Figure 1, we have proposed a ZIKV model that is calibrated on case data of GBS. ZIKV case numbers are obtained by scaling up from GBS. The advantage of this practice is that the GBS case numbers are more trustworthy and reliable compared to numbers obtained through surveillance of ZIKV where there is much scope for errors in the reporting rate. Our model considers heterogeneity in symptomatic and asymptomatic ZIKV infections (i.e., *θ* and *η*) as well as the local mosquito population (*m*). Model (1) was fitted to the reported excess GBS cases time series with different sets of parameters for symptomatic proportion (*θ*) and asymptomatic infectivity (*η*).

From a recent metadata study [20] and a serological study [19], the ratio *ρ* of GBS to symptomatic ZIKV cases was found to be 0.00024 and 0.00032 respectively (see Introduction of this study). Similarly, based on the data from eleven countries, Mier-y-Teran-Romero *et al.* [15] found the overall estimate for the risk of reported GBS “was 2.0 (95% CI 0.5-4.5) GBS reported cases per 10,000 ZIKV infections, i.e., 0.02%, (which is) close to the point estimate of 2.4 GBS cases per 10,000 ZIKV, i.e., 0.024 %, infections estimated using only data from French Polynesia”. In this study, the model estimation finds a ratio between GBS and symptomatic ZIKV cases as *ρ* = 0.000061 or equivalently *ρ* = 0.0061% with 95% CI 0.0050%-0.0086%. This or 1 GBS case per 16,393 ZIKV symptomatic cases which is approximately one quarter or 25% the magnitude of existing estimates. Our estimate, although still tentative and based on reasonable first approximations, seems plausible since ZIKV surveillance was generally unreliable and probably severely underreported, especially before 2016 [44, 52]. For this reason, we avoided using the ZIKV surveillance data to fit the epidemic model, and our estimate of *ρ* depends on the more reliable GBS data.

The model analysis estimated the IAR of ZIKV cases in NE Brazil to lie between 22% to 28% for the two waves. This is based on the assumption that the proportion of symptomatics *θ* = 0.5, which appears to be reliable according to the serological results of Aubry *et al.* [19]. This is in line with a number of model and empirical estimates for other areas of Brazil and South America. For example, Zhang *et al.* estimated some 18% IAR for the areas in Brazil [12]. In pointing this out, we must also note that most IAR estimates in the literature need to be treated with caution. Due to poor surveillance and limited knowledge about the ZIKV reporting ratio, the estimates may have been based on samples that are not representative of the general population as a whole.

Oliveira *et al.* [22] also identified a striking relationship between the dynamics in time of the first wave of excess GBS and that of microcephaly. Their Figure 1B shows the dynamics in time of these two conditions are almost identical apart from a delay of 23 weeks and differing otherwise by a scale factor. The remarkable similarity in the different epidemic time series allows us to compare the rates of GBS cases to those of microcephaly. By examining the peak heights of the two diseases, the ratio between them is 6.1 (maximum of microcephaly divided by maximum of GBS wave), which corresponds to 1 GBS case for every 6.1 microcephaly cases. If we make the reasonable assumption that the reporting rate of both conditions is similar, it is clear that GBS is a much rarer disease than microcephaly. Nevertheless, we still chose to predict ZIKV cases based on GBS rather than microcephaly cases, because of problems in the correct reporting of microcephaly over the study period. For example, the criteria for identifying microcephaly changed dramatically at different times over the two year period and in different areas, making the reporting coverage highly unstable. Moreover, previous to this period, reporting was not compulsory nor was there consistently defined criteria for identifying the condition.

Return now to the dynamics of the reconstructed ZIKV cases generated by Eqns (1) as calibrated on the GBS data (Fig 4). The reproductive number, ℛ_0_(*t*), which quantifies the transmission rate, was reconstructed by modelling the local meteorological data with Eqn (6). The estimated ℛ_0_(*t*) was found to oscillate due to seasonality between the values 1.1 < ℛ_0_ < 3.3, and on average was found ⟨ℛ_0_⟩ = 2.2. The average level and estimated range of ℛ_0_(*t*) are in line with previous studies [12, 44, 52]. Because of temperature dependence, ℛ_0_(*t*) reached minimum values in winters. The range of values the model predicted for ℛ_0_(*t*) is very similar to the intensities reported in Fig 3 of [12] for ZIKV in Brazil.

As the net growth rate of mosquitoes tends to increase as temperature increases [12, 54, 55], it is not surprising that our estimated *p*_1_ > 0 (the temperature dependence parameter in *m*(*t*)) is positive. The positive association between temperature and transmissibility has also been observed in the literature [52]. Significant nonzero estimates were found for parameters *p*_3_ and *p*_4_, which also control *m*(*t*), and thus the reproductive number ℛ_0_. This immediately suggests the existence of non-weather-driven temporal changes in the ZIKV transmissibility. The baseline drop in *m*(*t*) would also lead to a drop in ℛ_0_(*t*), and indicates a decrease in ZIKV transmissibility across the two epidemic waves. Since the official mandatory ZIKV reporting started on February 2016, this could have increased public awareness of ZIKV risk, and thus prevented infection effectively [53, 56, 57, 58, 59]. Disease control measures were also introduced by some local authorities during the second epidemic wave. The time-change point (*τ*_1_) when the baseline *p*_3_ switches to *p*_4_ in the model corresponds to March 7 of 2016. Interestingly, this time point coincides with the peak timing of the concurrent CHIKV outbreak [22]. Also, very close to this date, ℛ_0_(*t*) passed through a local minimum and then increased for a two month period, generating in turn an increase in GBS cases.

We compared the results of a non-mechanistic model in Supplementary Information S3 which did not take into account climatic factors, and those from our climate-driven model in Fig 4. Although the non-mechanistic model did not perform as well, it nevertheless provided useful insights by producing results that matched the impact of the daily temperature on ℛ_0_, the transmission of ZIKV.

Continuing further, we now attempt to estimate the reporting rate of ZIKV. We argue that the reporting rate of ZIKV disease increased dramatically around February and March of 2016, as suggested also in the literature [52]. Thus, it is reasonable to assume that the data for the second wave of ZIKV in 2016 is more reliable than that of the first. Taking the maximum of the second ZIKV wave divided by the maximum of the GBS wave, we find the ratio between the two diseases is 435.6; i.e., 1 GBS case per 435.6 reported ZIKV cases. However, our model fitting finds a ratio between GBS and symptomatic ZIKV cases as *ρ* = 0.000061, or 1 GBS case per 16,393 ZIKV symptomatic cases. Thus, we can conclude that the reporting ratio of symptomatic ZIKV cases is roughly 16393/435.6 ≈ 38. Namely for every 38 symptomatic ZIKV cases, there was 1 case reported, over the second wave in 2016. Hence we arrive at an estimate for the reporting ratio of ZIKV, namely 1:38. Moreover, as mentioned, when taking this reporting ratio into account our estimated IAR falls in the reasonable range 22% to 28% for the two waves.

Previous estimates of IAR relied on poor ZIKV data in Brazilian regions: some estimates appear to be less than 20%, and others yield more than 50% (see Supplementary Information S4). As mentioned, all these estimates must be treated with caution. This study is the first to use the more reliable GBS data as a proxy to estimate the IAR of ZIKV epidemics. We found that the IAR is likely to be below 33%.

In conclusion, we comment on the likelihood of a future major ZIKV outbreak in NE Brazil. Let us start from a “naive assumption” that the whole population (100%) in NE Brazil was susceptible to ZIKV at the beginning of 2015, even though it was probably less than 100%. Our results tell us that the estimated IAR is most likely below 33%. This indicates that after the 2015-2016 ZIKV outbreaks, probably more than (100 *-* 33% =) 67% of the population were susceptible and immune-naive. That is, *S*_*h*_ > 67%, after the last ZIKV outbreak that ended in 2016.

Recall that the effective reproduction number, ℛ_eff_ = *S*_*h*_ℛ_0_ < 1, must be less than unity to ensure the epidemic will not emerge. Given the susceptibility at the end of the outbreak was more than 67%, then we need ℛ_0_ < 1*/S*_*h*_ = 1/67% ≈ 1.5 to ensure ℛ_eff_ < 1 under the naive assumption, and no outbreak will emerge. An ℛ_0_ larger than approximately 1.5 will lead to a ZIKV outbreak. On the other hand, according to our estimation (Table 3), the initial susceptibility, *S*_*h*_(0), at the start of 2015 was likely to be below 75%. As Table 3 shows that, typically, *S*_*h*_(0) = 0.57% with 95% CI: 47% - 74%. Thus, at least (1 - 75% =) 25% of the (susceptible) population was not affected at all during the 2015-16 ZIKV epidemic waves in NE Brazil. It is possibly that this 25% (or more) of the population, were protected because of cross-protection from infection with other *Flaviviridae* and/or because of living in zones where ZIKV cannot persist, etc. With this possibility, we now have over ((100% *-* 25%) *-* IAR > (1 *-* 25%) *-* 33% =) 42% of the population who are still immune-naive and unprotected after the 2015-16 epidemic. Therefore, if ℛ_0_ < 1*/S*_*h*_ < 1/42% ≈ 2.38, a ZIKV outbreak will not occur. Now the estimated average ⟨ℛ _0_⟩ = 2.2 in Fig 4, which ensures ⟨ℛ_eff_⟩ < 1 and implies that it would be difficult for a ZIKV outbreak to appear in NE Brazil in the near future.

## List of abbreviations

CHIKV: Chikungunya virus
DENV: Dengue virus
ZIKV: Zika virus
GBS: Guillain-Barré syndrome
IAR: infection attack rate
BIC: Bayesian information criterion
NE: Northeastern
NB: negative-binomial
POMP: partially observed Markov process
CI: confidence interval
MLE: maximum likelihood estimate
PRCC: partial rank correlation coefficient

## Declarations

### Ethics approval and consent to participate

Since no personal data was collected, the ethical approval or individual consent is not applicable.

### Availability of data and materials

The epidemic time series data were kindly provided by Professor Oliveira from the Ministry of Health in Brazil, as used in their recent study [22].

### Consent for publication

Not applicable.

### Funding

The authors declare that this study is not funded.

#### Acknowledgements

The authors would like to acknowledge Professor Oliveira from the Ministry of Health in Brazil for kindly sharing the disease surveillance data used in this work.

### Disclaimer

The funding agencies had no role in the design and conduct of the study; collection, management, analysis, and interpretation of the data; preparation, review, or approval of the manuscript; or decision to submit the manuscript for publication.

### Authors’ Contributions

DH conceived the study. DH and SZ carried out the study. DH, SZ and LS discussed the results. DH, SZ, QL and LS drafted the first manuscript. DH, SZ, SSM and LS revised the manuscript. All authors gave final approval for publication.

## Supplementary Information

**S1 Case Definition of the Guillain-Barré Syndrome**

**S2 Brief Information of the Northeastern Brazil**

**S3 Fitting Results with Cubic Spline Reconstruction**

**S4 Brief Review of the Zika epidemics Infection Attack Rate**

